# Fabrication of radiopaque, drug-loaded resorbable polymer for medical device development

**DOI:** 10.1101/2025.01.30.635738

**Authors:** Sophie T. Melancon, Erin Marie San Valentin, Dominic Karl M. Bolinas, Marvin Bernardino, Archana Mishra, Gino Canlas, Gouthami Chintalapani, Megan C. Jacobsen, Allan John R. Barcena, Steven Y. Huang

**Affiliations:** Department of Interventional Radiology, The University of Texas MD Anderson Cancer Center, Houston, TX 77030, USA; College of Medicine, University of the Philippines Manila, Manila 1000, Philippines; Department of Chemistry, Lamar University, Beaumont, TX 77710, USA; Siemens Medical Solutions USA, Inc., Malvern, PA 19355, USA; Department of Imaging Physics, The University of Texas MD Anderson Cancer Center, Houston, TX 77030, USA

**Keywords:** Resorbable medical device, Gadolinium nanoparticles, Inferior vena cava filters, Radiopacity, Dipyridamole

## Abstract

Resorbable medical devices provide temporary functionality before degrading into safe byproducts. One application is absorbable inferior vena cava filters (IVCFs), which prevent pulmonary embolism (PE) in high-risk patients with contraindications to anticoagulants.

However, current absorbable IVCFs are limited by radiolucency and local clot formation risks. This study aimed to develop radiopaque, drug-loaded resorbable IVCFs with enhanced imaging and therapeutic capabilities.

Poly-p-dioxanone (PPDO) sutures were infused with gadolinium nanoparticles (GdNPs) and dipyridamole (DPA), an anti-thrombotic agent. GdNPs were synthesized with an average diameter of 35.76 ± 3.71 nm. Gd content was 371 ± 1.6 mg/g (PPDO-Gd) and 280 ± 0.3 mg/g (PPDO-Gd+DPA), while DPA content was 18.20 ± 5.38 mg/g (PPDO-DPA) and 12.91 ± 0.83 mg/g (PPDO-Gd+DPA). Suture thickness (0.39-0.49 cm, P = 0.0143) and melting temperature (103.61-105.90, P = 0.0002) statistically differed among the different groups, while load-at-break did not (4.39-5.38, P = 0.2367). Although suture thickness and melting temperatures differed significantly, load-at-break was preserved and did not alter the mechanical and degradation properties of the various IVCFs. Micro-CT imaging revealed enhanced radiopacity for Gd-containing IVCFs (2,713 ± 105 HU for PPDO-Gd, 1,516 ± 281 HU for PPDO-Gd+DPA). Radiopacity decreased gradually over 10–12 weeks. Clot-trapping efficacy was maintained, and no hemolysis or cellular toxicity was observed.

In conclusion, the GdNP- and DPA-infused PPDO IVCFs demonstrated improved radiopacity, anti-thrombotic potential, and compatibility with routine imaging, without compromising mechanical strength or safety.

## Introduction

Resorbable medical devices represent a significant advancement in biomedicine designed to perform critical functions such as structural support^1-3^ or drug delivery^4, 5^ for a temporary period. Typically, once their therapeutic purpose has been fulfilled, these devices gradually degrade into non-toxic byproducts that can be naturally absorbed or excreted by the body, eliminating the need for surgical removal. This feature not only reduces the risks associated with long-term implantation such as local infection^6^ or chronic inflammation,^7^ but also minimizes the need for follow-up surgeries, enhancing patient recovery and reducing healthcare costs.

As research continues to improve the performance and biocompatibility of resorbable materials, their potential extends to treating life-threatening conditions like pulmonary embolism (PE). PE affects nearly 900,000 people every year in the United States alone.^8^ Patients at risk of this condition typically receive anticoagulant therapy (e.g., low molecular weight heparin and direct-acting oral anticoagulants) to prevent the development of blood clots^8, 9^ and mitigate the risk of subsequent complications including strokes^9^ and cardiac arrest.^10^ However, for patients with contraindications to anticoagulant therapy, the implantation of inferior vena cava filters (IVCFs) is often typically recommended as an alternative treatment option.^11^ Traditionally, IVCFs are fabricated from metals such as nitinol or stainless steel and are intended for surgical retrieval after 25-54 days post-implantation as recommended by the Food and Drug Administration (FDA).^12^ Yet, only 17-30% of patients have these temporary filters removed despite the risk of complications such as filter migration and perforation.^13^ To address this issue, research has focused on developing resorbable IVCFs, which would eliminate the need for surgical removal.

One of the biggest challenges of making an effective resorbable IVCF is selecting a polymer that has the most optimal degradation time. It is critical that temporary IVCF remain fully functional for a minimum of 35 days following a major trauma or procedure, while also being removed prior to the onset of significant complications.^14^ Pre-clinical testing has been done to test the efficacy of absorbable IVCFs made with polylactic-co-glycolic acid (PLGA),^15^ polyglycolic acid (PGA),^16^ and poly-L-lactic acid (PLLA) and polycaprolactone.^17^ All three studies were successfully fabricated into filters with different degradation characteristics and clot trapping efficacies. On the other hand, Eggers et al.^18^ showed that poly-p-dioxanone (PPDO) is a promising polymer for fabricating absorbable IVCFs with a degradation time of 6 weeks.

PPDO is commercially available and inexpensive because of its common application as surgical sutures. However, unlike metallic IVCFs, PPDO filters cannot be easily visualized on conventional imaging such as computed tomography (CT) or X-ray. Visualizing the filter is critical for patient safety. It allows assessment of immediate and delayed filter complications, such as filter tilt, incomplete expansion, and device migration. Central venous access is obtained by passing a wire into the central veins. Being able to visualize the filter is critical to ensure that the catheter or wire used during the placement procedure does not displace the filter. Also, while IVC filters are designed to capture thrombi; the foreign body itself can also be thrombogenic. Depending on the severity of thrombus in situ below the filer, thrombectomy procedures could be performed which would require that the filter be visualized. Thus, for the safety of the patients, it is best that resorbable IVCFs be capable of being monitored with conventional x-ray imaging modalities. Previously, this has been addressed by the infusion of high atomic number (high-Z) NPs such as gold,^19^ bismuth,^20^ and tungsten.^21^

Gadolinium (Gd) is another high-Z element (Z=64) with attractive properties for enhancing imaging. Most commonly used as a contrast agent for magnetic resonance imaging (MRI), Gd chelates function by shortening T1- and T2-relaxation times, thus improving the clarity of MRI scans. Beyond MRI, Gd is also considered novel for its versatility in multiple imaging modalities such as CT imaging. In its NP form (GdNP), Gd offers additional advantages, including a significantly higher surface area, allowing the increased loading and therefore higher X-ray attenuation. This enhanced imaging capability and safety profile makes GdNPs particularly attractive for applications like infusion with PPDO IVCFs, as they can improve both the precision of filter placement and monitoring post-deployment outcomes through dual-modality imaging.

A significant challenge with most implantable medical devices, including IVCFs, is the high incidence of thrombosis. To address this issue, we hypothesize that coating IVCFs with antithrombotic drugs can reduce local clotting and mitigate this common complication.

Dipyridamole (DPA; 2,6-bis-diethanolamino-4,8-dipiperidinopyrimido(5,4-d)-pyrimidine), a drug known for its antiplatelet activity, shows promise in minimizing thrombosis when integrated into medical devices.^22^ This study aims to develop a novel absorbable IVCF by combining the resorbable polymer PPDO with radiopaque gadolinium nanoparticles (GdNPs) and the antithrombotic drug DPA. The fabricated filters were evaluated for their physicochemical properties and degradation behavior over time in an in vitro system. This work represents the first preclinical development of an absorbable IVCF that integrates both imaging capability and therapeutic functionality.

## Materials and Methods

### Synthesis and Characterization of Gadolinium Nanoparticles

GdNPs were synthesized by heating 5 mmol gadolinium acetate (ThermoFisher Scientific, Waltham, MA) at 120°C for 1 h, then at 320°C for 6 h in the presence of stabilizers oleic acid (technical grade, 90%), oleylamine (technical grade, 70%), and 1-octadecene (technical grade, 90%) under argon gas protection. The resulting metallic white solution was washed with ethanol, centrifuged, and redispersed in dichloromethane (DCM; ACS reagent grade, ≥99.5%, ThermoFisher Scientific) until use.

The size and morphology of synthesized GdNPs were visualized and characterized by transmission electron microscopy (TEM) using a JEM 1010 microscope (JEOL USA, Peabody, MA) and the size and distribution of the nanoparticles were estimated using ImageJ software (National Institutes of Health, Bethesda, Maryland).

### Fabrication of radiopaque, drug-loaded resorbable IVCFs

IVCFs were braided by modifying an agglomerate cork (32.6 mm dimeter x 36.6 mm height) with 15 zinc finishing nails (Everbilt #19 x ½ in) placed equidistantly (2.17 mm apart) at the top and bottom circumferences of the cork. WebMax 2-0 poly-p-dioxanone sutures (PPDO) (Patterson Veterinary, Liberty, MO) were wrapped around the cork by alternately looping through a nail on top and then at the bottom with 8 nail intervals. After every two turns, the suture goes up and through a round 3D-printed resorbable tip (polypropylene, 6.2 mm diameter x 1.2 mm height), printed using an S7 Ultimaker 3D-printer and designed using UltiMaker Cura 5.5.0. This forms the conical portion of the IVCF. After braiding, the filters were annealed using an oven at 150 °C. The resulting filters measured 60 mm (length) × 30 mm (diameter) and included a stent portion and a conical portion (**Supplementary Figure S1**).

Prior to coating with NPs, the IVCFs were soaked in DCM to wash off the purple coating. Filters were infused with either GdNP only (PPDO-Gd), DPA only (PPDO-DPA), or Gd-DPA (PPDO-Gd+DPA) solution containing 2.5 % polycaprolactone (PCL; Sigma-Aldrich, St. Louis, MO) using a wet dipping method. Coated sutures were characterized using micro-CT (Skyscan 1276; Bruker, Billerica, MA), field-emission scanning electron microscope (Nova NanoSEM 230; FEI, Hillsboro, OR), energy-dispersive x-ray spectroscopy (EDAX Element EDS; Ametek, Berwyn, PA) and analyzed using TEAM WDS Analysis System software (version V4.5.1-RC13 (Ametek). The mechanical strength of sutures was also measured using an Expert 7601 Tension Testing System (ADMET, Norwood, MA). Calculation of the % Gd/DPA remaining was obtained by subtracting the amount released from the total amount of Gd/DPA in the IVCF divided by the total Gd/DPA, multiplied by 100. Total Gd/DPA was determined by adding the Gd/DPA released plus the Gd/DPA remaining in the IVCF. The total Gd/DPA content was calculated by taking the total Gd/DPA and dividing with the weight of the IVCF, multiplied by 100.

### Radiopacity over time and in vitro Gd and DPA release studies

To monitor the release of GdNP and DPA in vitro, the stents were soaked in 20 mL phosphate-buffered saline (PBS) at pH 7.4 and 37 °C. The PBS solution was collected and changed at days 1, 3, 5, and weekly until 12 weeks. DPA in solution was quantified at 295 nm using a UV-Vis spectrophotometer (Cary WinUV 60 SW, Agilent Technologies, Stanta Clara, CA). Gd solutions were prepared for elemental analysis by dissolving 1 mL aliquot with 1 mL HNO3, and shaken at room temperature overnight. Samples were then diluted to 15 mL with 2% HNO3 for quantification of Gd in an atomic emission spectrometer (Agilent MP-AES; Agilent Technologies, Santa Clara, CA), using Gd as a standard for calibration.

### Biocompatibility of the materials

Cytotoxic effects of GdNP were tested against mouse vascular aortic smooth muscle cells (MOVAS) and immortalized human vascular endothelial cells (EC-RF24). Cells were grown in 96-well plates with complete media for 24 h before incubating with GdNP, DPA, and Gd-DPA-treated sutures at varying concentrations. Cell viability was measured after 24 h with alamarBlue cell viability assay (ThermoFisher Scientific, Waltham, MA).

Hemolysis assay was performed by modifying the protocol by Evans et al. (2013). Erythrocytes were collected by placing fresh rat blood in green-top heparinized vacutainers (BD Vacutainer, Franklin Lakes, NJ) were washed with cold PBS and centrifuged to collect erythrocytes. Positive controls were added with 50uL of 10% Triton X-100, while negative controls were added with 50 μL of PBS. Remaining wells were added with three sterilized 1 cm strands of the control PPDO, PPDO-Gd, PPDO-DPA or PPDO-Gd+DPA. Percent lysis was calculated using the optical densities (OD) of the samples against samples containing Triton-X 100 solution. Samples were analyzed in triplicates and percent (%) hemolysis was calculated by subtracting the absorbance of the PBS control from the absorbance of the sample, dividing the difference by the difference between the absorbances of the positive and negative controls.

### *In vitro* clot trapping efficacy experiment

First, alginate-based clots were synthesized using 2 % sodium alginate solution and iron nanoparticles (0.1 mg/mL)/fluorescent dye rhodamine B (0.1 mg/mL). Once thoroughly homogenized, 1 mL suspension solution was mixed with 100 mM CaCl2 solution in a 2 ml Eppendorf tube and allowed to solidify overnight at room temperature. Solidified clots were washed with milliQ water, cut (5 mm x 5 mm x 20 mm) and stored at 4°C until further application.

A silicone vena cava model (United Biologics, Irvine, CA) connected to a Flowtek 125 pump (United Biologics, Irvine, CA) was used to study clot-trapping efficacy of the fabricated IVCFs. IVCFs were first implanted by inserting it through a designated insertion pouch. This was securely fastened using a screw clamp before setting the flow pump. The pump was then filled with about 1 L of water and allowed to flow through the model while removing air bubbles. To ensure constant flow of water throughout the model, the pump was set at a continuous pulse at 40% flow rate.

Clots (5 mm x 5 mm x 20 mm) were deployed through the right posterior access site in the model (representing the right iliac vein) which is secured with a hemostatic valve. The ratio of maximum diameter of clot to filter is 0.33. The clots were allowed to flow towards the IVCF and clot trapping efficiency was evaluated whether the clots were captured by the filters or not. Each cylindrical clot was only introduced once in the flow pump.

Clot trapping efficiency experiments were also observed under fluoroscopy using a Siemens SOMATOM Definition Edge scanner (Siemens Healthineers, Erlangen, Germany). To replicate soft tissue in an animal or human model, saline bags were placed over the IVC silicone model.

### Statistical Analysis

All studies were conducted in triplicate (*n* =3) or three independent experiments. The data were presented as mean ± SD. GraphPad Prism, version 10.3.1 (464) software (GraphPad, San Diego, CA) was used to perform all statistical analyses. A two-tailed t-test or one-way analysis of variance (ANOVA) was used where appropriate, and differences were considered significant at P*<*0.05.

## Results

### Synthesis and characterization of the GdNP

The one-pot solvothermal technique of synthesizing GdNPs allowed for uniform sizing and shape of the nanoparticles as seen on the TEM in **Figure 1A**. The nanoparticles were seen to have a plate-like structures with an average diameter of 35.76 ± 3.71 nm (**Figure 1B**).

**Figure 1.**
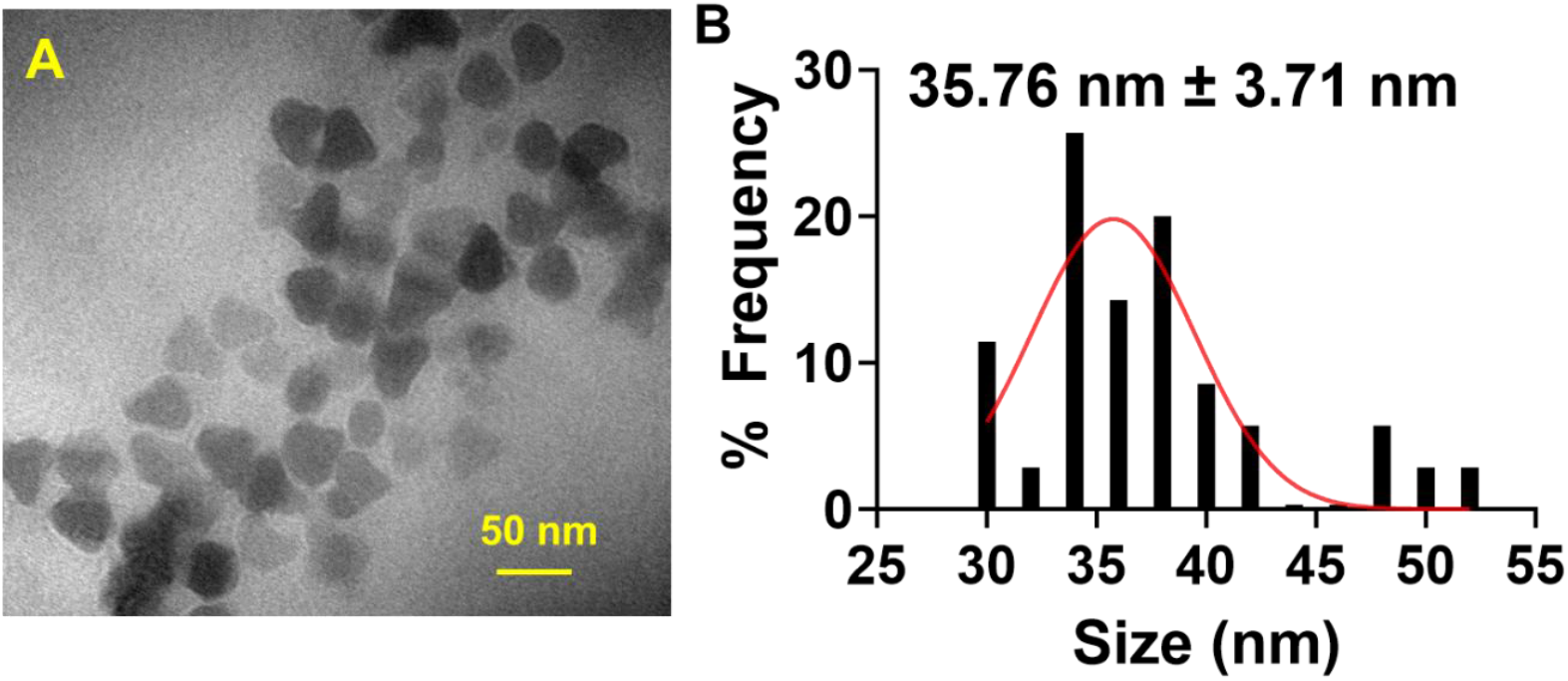
Synthesis and characterization of gadolinium nanoparticles (GdNPs). (A) Transmission Electron Microscopy (TEM) image shows that the synthesized GdNPs have a plate-like structure. (B) Quantification of the size of GdNPs using ImageJ software resulted in an average diameter of 35.76 ± 3.71 nm.

### Fabrication of the drug-loaded, radiopaque, and resorbable IVCFs

Synthesized GdNPs were incorporated with PCL solution in preparation for infusion onto the PPDO sutures. Compared to the bare PPDO (control), the GdNP-, DPA-, GdNP+DPA-infused PPDO sutures resulted in a rough surface as seen on SEM (**Figure 2, top**). The presence of GdNPs at 1.185 keV were also confirmed using SEM-EDX (**Figure 2, bottom**).

**Figure 2.**
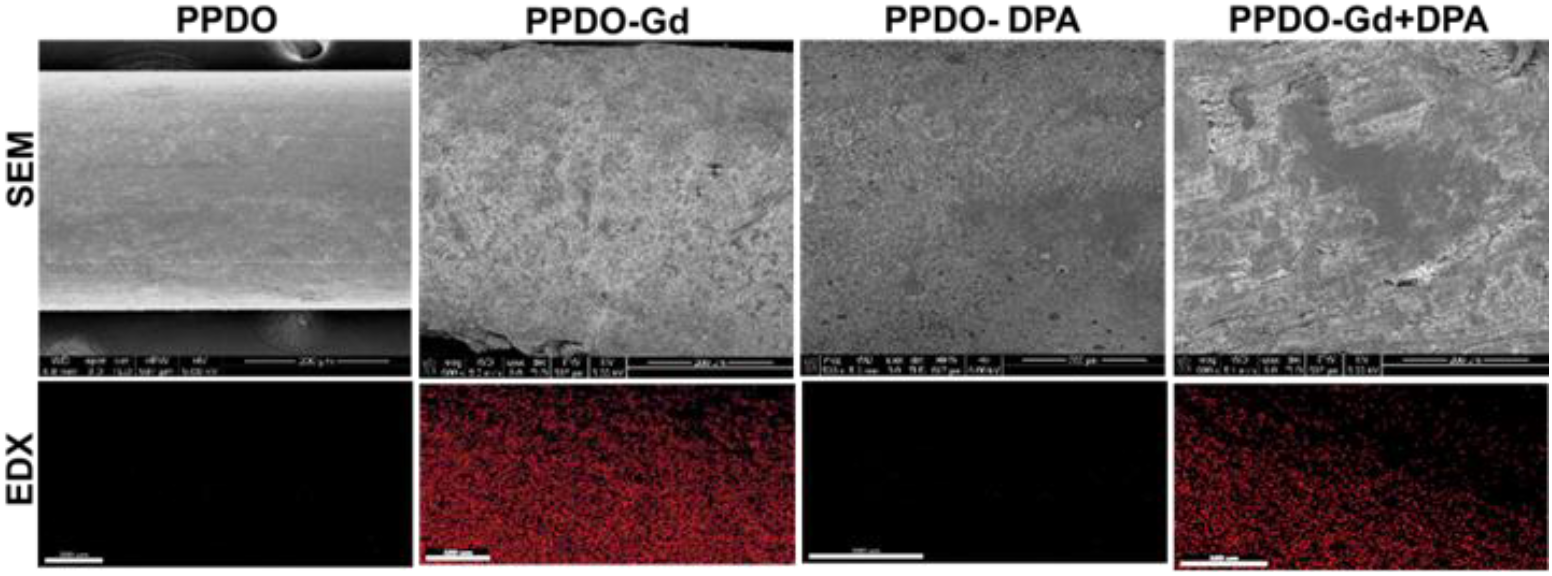
Scanning electron microscopy (SEM) and gadolinium (Gd) mapping using energy-dispersive X-ray (EDX) spectroscopy. Control poly-p-dioxanone (PPDO) sutures had a smooth surface, and addition of GdNPs increased the roughness of the surface of the PPDO. EDX confirmed the presence of Gd in PPDO-Gd and PPDO-Gd+DPA.

The method of braiding the IVCFs using a makeshift base and annealing proved effective as seen in **Figure 3**, wherein the devices were able to hold their shape even after being removed from the base. The commercially available PPDO suture is dyed purple to improve its visibility during surgical procedures, as illustrated in Figure 3. This purple dye dissolves upon exposure to DCM. The infusion of GdNP, which is white, turns the suture white (PPDO-Gd), while the addition of yellow DPA imparts a yellow coloration to both PPDO-Gd and PPDO-Gd+DPA. This shows the successful infusion of GdNP and/or DPA within the PPDO suture. Initial X-ray and mCT imaging show that the PPDO-Gd and PPDO-Gd+DPA had increased radiopacity as compared to the control PPDO and PPDO-DPA alone.

**Figure 3.**
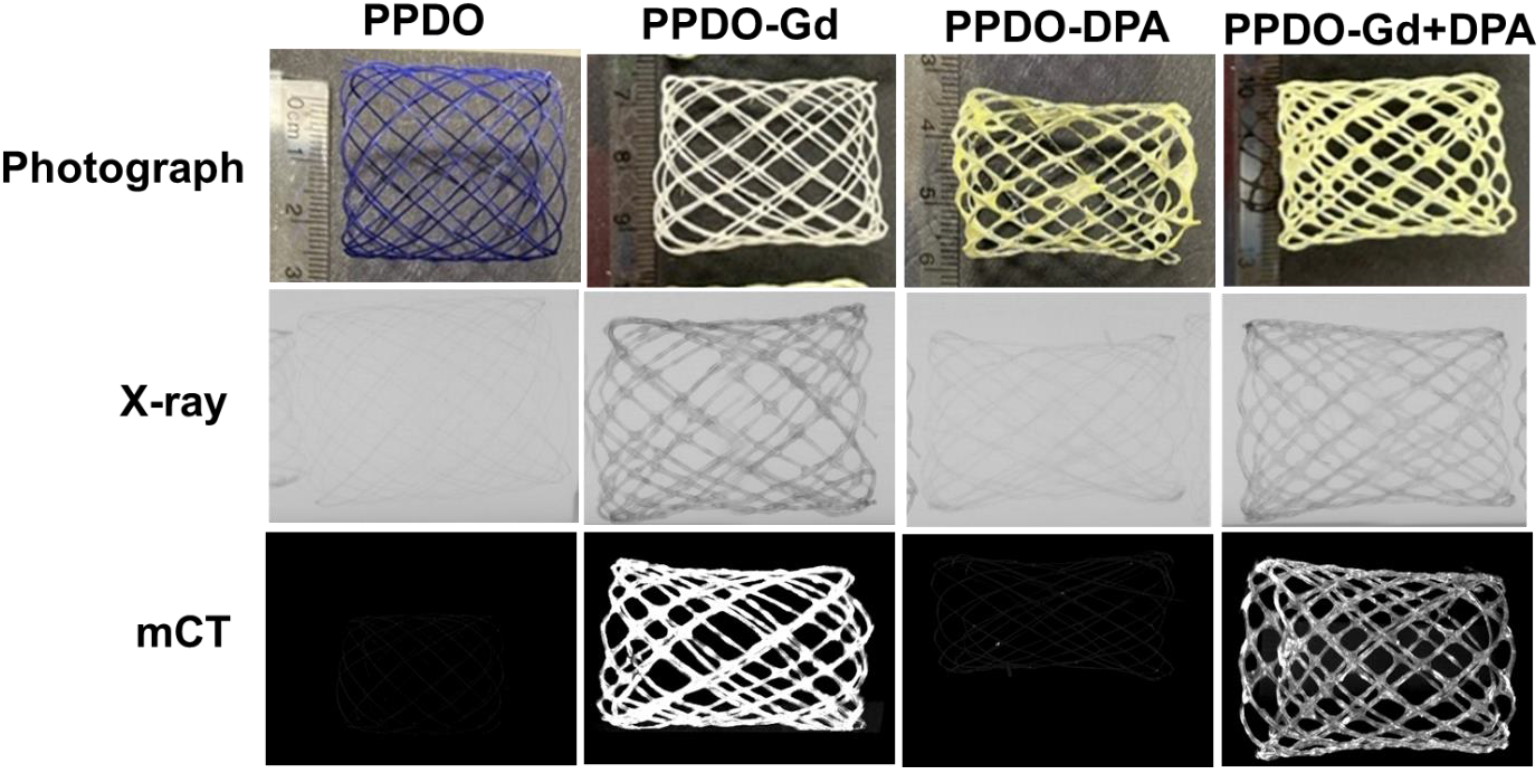
Comparison of the photograph, X-ray, and micro-computed tomography (mCT) images of the various poly-p-dioxanone (PPDO) formulations. Gadolinium (Gd)-containing PPDO sutures (PPDO-Gd and PPDO-Gd+DPA) had increased signal enhancement in X-ray and mCT imaging as compared to control PPDO and PPDO-DPA. DPA, dipyridamole.

### Physicochemical characterization

Table 1. shows the baseline physicochemical characteristics of the distinct formulations. Suture thickness (P = 0.0143) and melting temperature (P = 0.0002) statistically differed among the different groups, while load-at-break did not (P = 0.2367). Gd content was statistically higher in PPDO-Gd (P < 0.0001) compared to PPDO-Gd+DPA, while bare PPDO and PPDO-DPA did not have any trace Gd detected in elemental analyses. These also corresponded well with the radiopacities measured using mCT, where radiopacity of PPDO-Gd was also statistically higher (P = 0.0023) compared to the radiopacity of PPDO-Gd+DPA.

**Table 1.** Physicochemical properties of the inferior vena cava filters.

| | Suture thickness (cm) | Melting temperature, $T_m$ ($^{\circ}\text{C}$) | Load-at-break (kg) | Gd content (mg/g) | Radiopacity (HU) | DPA content (mg/g) |
| --- | --- | --- | --- | --- | --- | --- |
| **Control** | $0.39 \pm 0.00$ | $105.90 \pm 0.30$ | $5.24 \pm 0.12$ | 0 | $-130 \pm 38$ | 0 |
| **PPDO-Gd** | $0.49 \pm 0.05$ | $103.32 \pm 0.68$ | $4.39 \pm 0.87$ | $371 \pm 1.60$ | $2,713 \pm 105$ | 0 |
| **PPDO-DPA** | $0.41 \pm 0.00$ | $104.13 \pm 0.20$ | $4.60 \pm 0.64$ | 0 | $-135 \pm 172$ | $18.20 \pm 5.38$ |
| **PPDO-Gd+DPA** | $0.42 \pm 0.03$ | $103.61 \pm 0.08$ | $5.38 \pm 0.65$ | $280 \pm 0.31$ | $1,516 \pm 281$ | $12.91 \pm 0.83$ |

Abbreviations: DPA, dipyridamole; Gd, gadolinium; HU, Hounsfield unit; PPDO, poly-p-dioxanone.

### Radiopacity over time and Gd and DPA release studies

**Figure 4A** illustrates the radiopacity of the PPDO, PPDO-Gd, PPDO-DPA, and PPDO-Gd+DPA sutures over time. It is evident that the Gd-containing IVCFs had high HU values which gradually decreased over time. It can also be noted that PPDO-Gd had almost 2x higher values than that of PPDO-Gd+DPA. At Week 10, about 10% of GdNPs remain in the sutures (**Figure 4B**) as estimated using elemental analysis. Along with radiopacity, the load-at-break also slightly decreases over time with no significant differences among the various PPDO formulations (**Figure 4C**). DPA release shows a similar trend as that of the Gd release, where there is an initial decrease of about 20% followed by slow steady release within the 6-week period and a sharp decline at 8 weeks for DPA-containing IVCFs revealed (**Figure 4D**).

**Figure 4.**
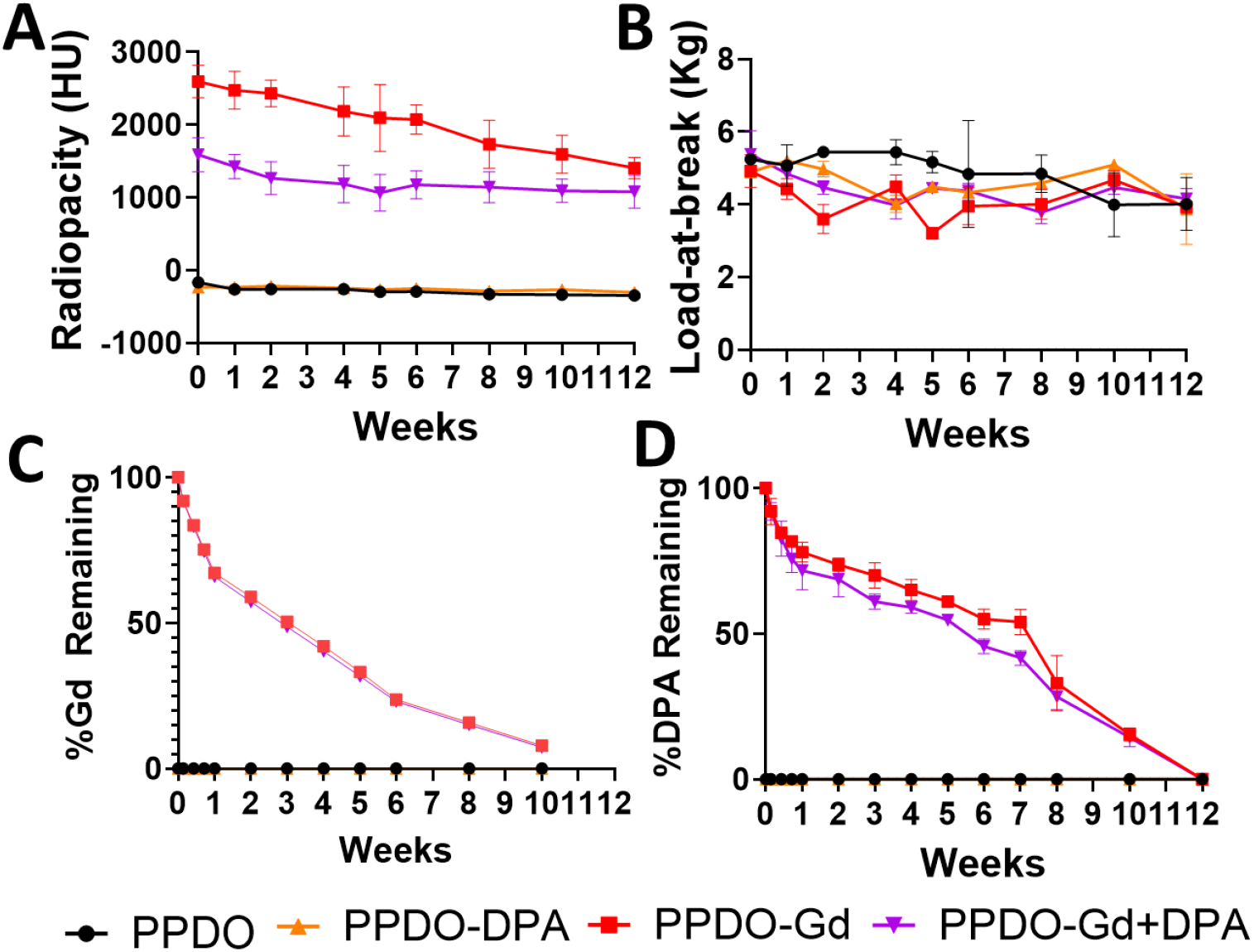
Longitudinal monitoring of radiopacity, mechanical strength, gadolinium (Gd) and dipyridamole (DPA) release over 10-12 weeks. (A) Radiopacity, measured in Hounsfield units (HU), decreased gradually over time, mirroring the decreasing trend in mechanical strength (B), assessed as load-at-break in kilograms (kg). (C, D) Gd and DPA content showed steady reductions, reflecting their controlled release over the study period. PPDO, poly-p-dioxanone.

### *In vitro* hemolysis and cell viability assays

Biocompatibility of the GdNP- and DPA-infused IVCFs were tested against RF24 (**Figure 5A**) and MOVAS (**Figure 5B**) cells. In both cell lines, the cells grown in different concentrations of treated media did not induce significant reduction in cell viability. In addition, the hemolysis assay using rat red blood cells indicated that only Triton X-100 (positive control) showed significant hemolysis across all treatment groups (**Figure 5A**).

**Figure 5.**
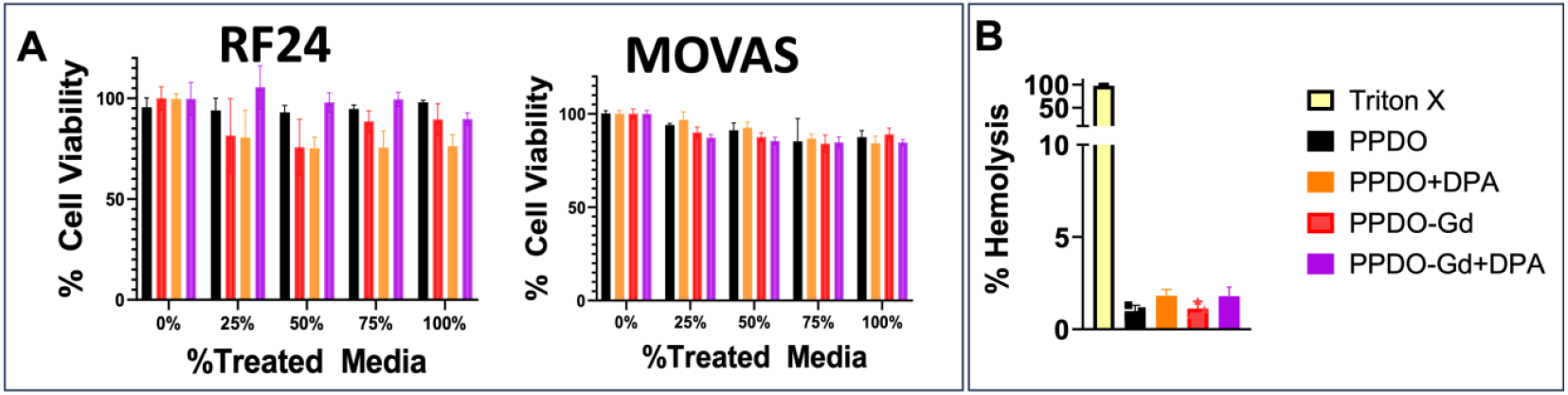
In vitro cell viability and hemolysis assays. (A) Treatment of immortalized human vascular endothelial (EC-RF24) and smooth muscle cells (MOVAS) with varying concentrations of treated cell media with all formulations did not show significant cytotoxicity (P <0.05). Cells were incubated in treated media for 24h before cytotoxicity assay with 10% alamarBlue. (B) Hemolysis assay shows no significant differences among the various groups. Triton X was used as the positive control. DPA, dipyridamole; Gd, gadolinium; PPDO, poly-p-dioxanone.

### Assessing the clot-trapping efficiency and imaging of IVCFs

The clot-trapping efficacy of the fabricated absorbable IVCF was evaluated using a commercial micro-circulation system designed to mimic human venous flow (**Figure 6A, left**). The system, filled with ∼1 L of water, was set to operate at a continuous pulse with a 40% flow rate to simulate physiological conditions. The IVCF was placed inside the device, after which an artificial alginate clot (5 × 5 × 20 mm), dyed pink for enhanced visibility, was introduced into the flow model (**Figure 6A, right**).

**Figure 6.**
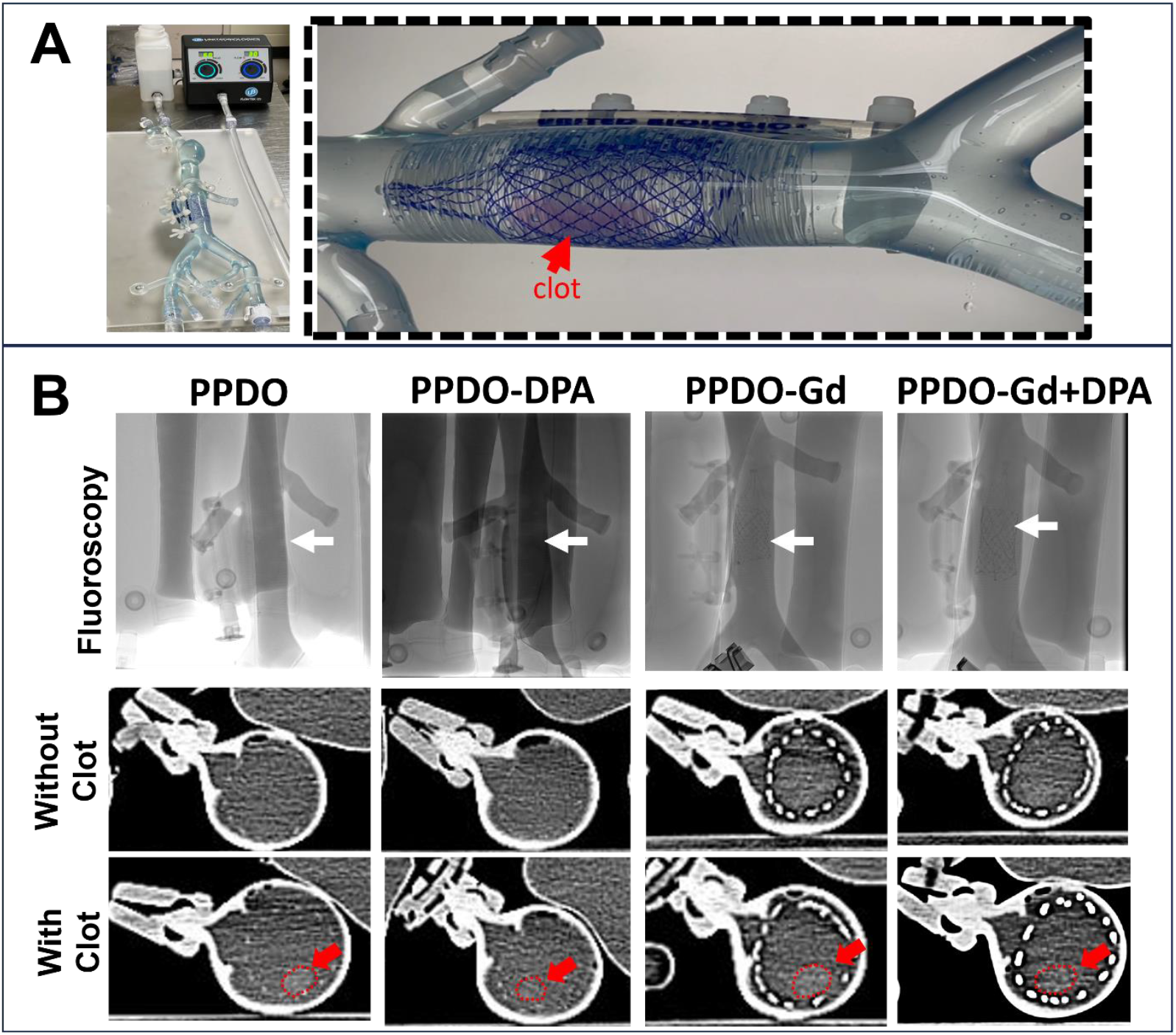
Clot trapping efficacy and imaging. A vena cava flow model (A, left) was used to evaluate the in vitro clot trapping of the various IVCFs at week 0 using an artificial alginate clot (5 mm × 5 mm × 20 mm). The system contained ∼1 L of water at 40% flow rate. (A, right) All the fabricated resorbable inferior vena cava filters (IVCFs) successfully captured the clot (red arrow, circled). X-ray imaging of the system included fluoroscopy (top) and CT with and without clots (bottom), demonstrating the enhanced visibility of the IVCF (B). White arrows indicate position of the IVCF in fluoroscopy where only gadolinium (Gd)-containing IVCFs are visible. Strands of the Gd-containing IVCF, including PPDO-Gd and PPDO-Gd+DPA, exhibited clear radiopacity under both imaging modalities. Clots are difficult to see with standard X-ray imaging. DPA, dipyridamole; PPDO, poly-p-dioxanone.

The fabricated IVCF exhibited successful clot-trapping capabilities, effectively securing the clot within the filter’s structure. Imaging was performed using fluoroscopy and micro-CT (mCT) to assess radiopacity and the filter’s visibility in situ. Under fluoroscopy (**Figure 6B, top**), the gadolinium (Gd)-containing IVCF showed excellent radiopacity, with individual strands clearly visible. Micro-CT imaging (**Figure 6B, bottom**) provided enhanced visualization of the system, showing the filter both with and without the captured clot. Red arrows highlight the position of the clot (encircled), which is not visible under standard X-ray-based imaging systems.

## Discussion

Temporary medical devices are critical in providing short-term support to patients with various medical conditions, with the intent of removal once the condition improves or resolves. A significant advantage of using absorbable materials, such as resorbable polymers, is that they can provide the necessary functionality during their active period and then gradually dissolve, being safely absorbed by the body, thus eliminating the need for surgical extraction. This approach not only minimizes the risks associated with additional surgeries but also enhances patient comfort and recovery. In this study, we introduced a multifunctional, polymeric IVCF fabricated from PPDO, which is not only absorbable but also imageable, incorporating GdNPs for radiopacity and serving as a depot for DPA to reduce clot formation. This combination allows real-time imaging and controlled drug delivery, potentially improving the therapeutic efficacy and safety of temporary implants.

GdNPs have been previously shown as an imaging enhancer for various medical devices.^23, 24^ Although more known as an MRI agent due to its paramagnetic properties, gadolinium is also a heavy metal like iodine which attenuates X-rays.^25^ With a higher atomic number (Z = 64) compared to iodine (Z = 53), gadolinium has a stronger ability to absorb X-ray energy. It also features a higher K-edge energy (50 keV), which is better aligned with the peak intensity of the X-ray spectrum generated during CT scans, typically ranging from 50–60 keV at 80–140 kVp settings in modern CT systems. As a result, gadolinium effectively absorbs more of the X-ray energy, making it a more efficient attenuator during clinical CT imaging.^25^

To efficiently incorporate gadolinium into absorbable polymers, such as PPDO, we utilized a nanoparticle form of gadolinium and employed a wet-dipping method alternating with PCL. GdNPs have a high surface area that allows them to easily integrate into the polymeric matrix of PPDO. The addition of PCL enhances the surface roughness of PPDO, improving the entrapment of gadolinium within the polymer, as seen in the studies of Damasco et al.^20^ and San Valentin et al.^21^ Our study confirmed that adding PCL to PPDO increased the surface roughness, facilitating greater gadolinium infusion, as observed in the SEM images (Figure 2). This modification also enabled sustained release of gadolinium over time, with 7.88% remaining in PPDO-Gd and 7.36% in PPDO-Gd+DPA at Week 12 (Figure 4A). However, the release of gadolinium did not correspond directly to a decrease in radiopacity. Even with approximately 7% of gadolinium remaining, the radiopacity was still intense. For instance, the radiopacity of PPDO-Gd decreased from 2594 ± 222 HU at Week 0 to 1406 ± 147 HU at Week 12 (approximately 46% reduction), and for PPDO-Gd+DPA, it decreased from 1589 ± 233 HU at Week 0 to 1078 ± 227 HU at Week 12 (approximately 32% reduction). This discrepancy may be due to the high initial radiopacity, meaning that even with a relatively small amount of gadolinium remaining, it still produced a strong signal in micro-CT imaging. Fine-tuning the concentration to start with less Gd and lower radiodensity could allow more precise monitoring of the Gd levels.

Beyond GdNPs, this study also focused on DPA, an antiplatelet drug known to inhibit platelet aggregation and prevent thrombosis. DPA’s incorporation into the IVCF offers potential for enhancing the filter’s antithrombotic properties, minimizing clot formation and improving device efficacy. Previous studies^26^ have demonstrated that DPA-loaded devices resist platelet adhesion without compromising hemocompatibility, suggesting that its inclusion could further improve the IVCF’s ability to prevent thrombosis. DPA works by inhibiting platelet aggregation, thus preventing the formation of blood clots, which is vital for minimizing the risk of thrombosis in the IVCF. By reducing platelet activation, DPA enhances the overall anti-thrombotic properties of the filter, promoting its effectiveness in capturing and preventing the propagation of thrombi. Similarly, Dominguez-Robles, et al.^27^ developed a 3D-printed drug-loaded cardiovascular grafts wherein they showed resistance of the device to platelet adhesion without affecting hemocompatibility and cytocompatibility in HUVEC cells. In addition, their group showed that an increased loading of DPA onto the device also corresponded to a more effective antiplatelet activity, suggesting that our study can be further improved by exploring different concentrations of loaded DPA. Several studies on drug-coated (e.g. rapamycin, heparin, and paclitaxel) metallic IVCFs have been explored primarily to combat neointimal hyperplasia, ^28, 29^ but this is the first study on DPA-loaded absorbable IVCFs. The systemic effect as the DPA resorbs was not evaluated during this study. We plan to assess systemic concentrations of DPA in future studies.

Although there were no significant differences load-at-break among the groups tested at week 0, the thickness and melting temperature (Tm) of PPDO slightly decreased with the addition of GdNP and DPA. This decrease in Tm is likely due to modifications in the crystalline and amorphous regions of PPDO caused by the incorporation of GdNPs and DPA, a phenomenon observed in previous studies of polymer composites.^30, 31^ Despite these initial differences in Tm, our in vitro evaluation demonstrated that all fabricated IVCFs, including those loaded with GdNPs and DPA, degraded over time without negatively affecting the material’s inherent degradation properties. Although we expected to have a sharp decrease in load-at-break starting at week 6 as shown by Egger, et al.^32^, we did not see this trend in all of our IVCFs. This is due to the in vitro system that was used in the study where IVCFs were soaked in PBS at room temperature with shaking compared to Eggers’ system that used a more elaborate closed-loop venous flow simulator with controlled flow, pressure, pH, temperature, viscosity and density to mimic blood.^18, 32^ On the other hand, the release profiles of GdNPs and DPA followed a predictable trend, with a rapid initial release phase transitioning into a steady release phase, consistent with the degradation kinetics observed for radiopaque resorbable IVCFs previously developed.^20, 21, 33^

Additionally, the IVCFs demonstrated excellent biocompatibility in vitro, as evidenced by the lack of cytotoxic effects and the maintenance of cell viability in endothelial and smooth muscle cell lines. One of the most significant advantages of this approach is the clot-trapping capability of the IVCF, which was demonstrated in a controlled vena cava flow model. The 40% flow rate in a vena cava flow model represents a controlled percentage of the system’s maximum flow capacity, used to simulate blood flow under reproducible, physiological conditions. In the human inferior vena cava (IVC), the average blood flow rate is typically around 2 to 3 liters per minute. Setting the flow rate to 40% of the system’s maximum capacity mimics this normal blood flow at approximately 1.2 liters per minute, providing a moderate yet accurate representation of IVC dynamics.^34^ This flow rate ensures that the experiment avoids excessive turbulence while still reflecting realistic blood flow, allowing for effective testing of IVC filters and other medical devices in a controlled environment. We opted for alginate because of their ability to mimic the mechanical properties of blood clots, providing a stable and reproducible model for testing medical devices.^35, 36^ Alginate’s gel-like properties allow for precise control over clot size and consistency, ensuring standardized experimental conditions. It is non-biological, reducing safety concerns, and is easy to handle in laboratory settings. Additionally, alginate clots can be dyed to improve visibility under imaging techniques and may also be loaded with imaging agents for fluoroscopy and CT. Results in Figure 6B shows the enhanced visibility of the IVCFs containing Gd (PPDO-Gd and PPDO-Gd+DPA), even the single strands using the axial micro-CT images. Same is true for fluoroscopy images which are helpful for device implantation since usually, this is the imaging modality that is used when the device is placed.

Although the results from this study are promising, there are several limitations. First, the in vitro degradation testing may not fully replicate in vivo conditions, where biological factors, such as blood flow and pressure, pH, and presence of enzymes, could influence the degradation rate and performance. Second, the absence of in vitro and vivo testing for DPA’s efficacy and long-term biological effects leaves some uncertainties regarding its performance in a living organism. Future studies involving large animal models are needed to further assess the degradation, efficacy, and overall performance of the PPDO-Gd-DPA IVCF in a dynamic biological environment. Additionally, the design and findings from this study could be translated to other absorbable medical devices, such as stents and sutures, with temporary functionality and long-term risks, offering potential for improvements in patient care across multiple clinical areas.

## Conclusion

This study demonstrates the feasibility of incorporating GdNPs and DPA into absorbable IVCFs to enhance both imaging capabilities and therapeutic functionality. The GdNPs successfully improved the radiopacity of the filter, enabling better visualization under multiple imaging modalities, while the potential of DPA to reduce thrombosis presents an exciting avenue for future research and applications for IVCFs. However, limitations such as the lack of *in vivo* testing for this novel device and the limited clot-trapping duration of the filter due to its absorbable nature highlight the need for further investigation. Furthermore, large animal models are essential to validate the long-term efficacy and safety of this novel approach. Overall, the development of absorbable IVCFs with dual imaging and therapeutic properties offers promising potential not only for vascular interventions but also for other medical applications requiring temporary yet effective treatment solutions.

## Acknowledgments

The authors would also like to acknowledge, K. Dunner at the High-Resolution Electron Microscopy Facility for assisting with the transmission electron microscopy and Dr. James Gu from the Houston Methodist Electron Microscopy Core for his expertise in SEM-EDX mapping.

## Data sharing statement

The data that support the findings of this study are provided as supplementary materials.

