## Supplementary Figure S1 for "Fabrication of radiopaque, drug-loaded resorbable polymer for medical device development"

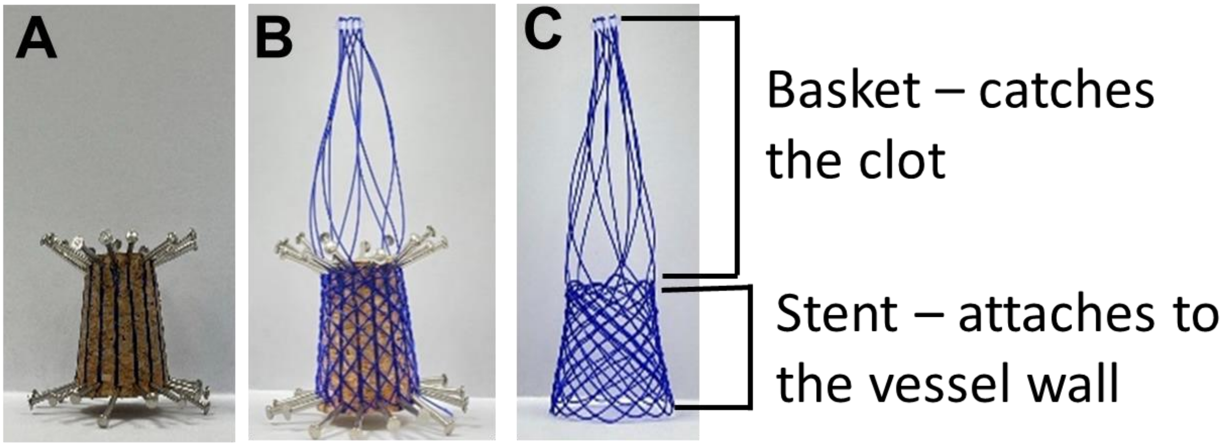


**Supplementary Figure S1. Fabrication process of the inferior vena cava filter (IVCF).** (A) Nails were evenly spaced and secured to a cork as a guide for braiding. (B) Poly-p-dioxanone (PPDO) sutures were braided around the nails and a 3D-printed plastic tip to form the IVCF structure. (C) After braiding, the nails were carefully removed, leaving behind the braided IVCF. The IVCF consists of two main components: the basket, which captures clots, and the stents, which anchor the device to the vessel wall for stability and function.
